# Senolytic treatment guided by carotid stenosis confers protection against acute ischemic stroke in aged rodents

**DOI:** 10.64898/2026.04.24.720745

**Authors:** Szilvia Kecskés, Péter Makra, Tamás Monostori, Zoltán Veréb, Ferenc Bari, Ákos Menyhárt, Eszter Farkas

## Abstract

**Background:** Advanced age is associated with larger infarct volumes and poorer functional recovery after acute ischemic stroke (AIS). Carotid stenosis is also a common comorbidity in older individuals and often predicts subsequent AIS. However, no age-specific therapy is currently available to protect the aging brain from aggravated ischemic injury. Here, we investigated whether a senolytic approach could improve cerebrovascular status and reduce ischemic brain injury in a comorbid aging model of AIS.

**Methods:** Unilateral common carotid artery occlusion was induced in young and aged rats and served as a diagnostic trigger for chronic senolytic therapy with dasatinib plus quercetin (D+Q). Two weeks later, the distal middle cerebral artery was occluded for 60 min. Compared with untreated animals, infarct size was measured, spreading depolarizations (SDs) were recorded electrophysiologically, cerebral blood flow (CBF) dynamics were monitored by laser speckle contrast imaging, and cerebrovascular senescent cell burden was assessed by immunocytochemistry. Cerebral angiogenesis, central and systemic inflammatory markers, and metabolic status were evaluated using protein arrays and blood glucose measurements.

**Results:** Aged rats developed larger infarcts than young controls, and this age-related increase was attenuated by D+Q treatment. D+Q reduced the higher frequency of SDs observed in the aged ischemic brain. Increased cerebrovascular senescence in aged animals was diminished by D+Q, accompanied by enhanced angiogenesis, although CBF responses to SDs and reperfusion were unchanged. In addition, D+Q modulated central and systemic inflammatory profiles and counteracted age-related metabolic impairment.

**Conclusions:** Senolytic D+Q therapy administered after carotid artery occlusion confers multifaceted protection against subsequent AIS in the aged brain. By targeting fundamental aging mechanisms that exacerbate brain vulnerability to AIS, D+Q enhances the resilience of the aging neurovascular niche. These results identify senolytic therapy as a promising preventive personalized approach to mitigate the disproportionate impact of AIS in older individuals and warrant further investigation.

## Background

Aging is a dominant biological determinant of acute ischemic stroke (AIS), rather than merely a demographic risk factor. Advanced age is associated with larger infarct volumes (Ay et al., 2005), impaired functional recovery (Knoflach et al., 2012), and diminished responsiveness to reperfusion and neuroprotective therapies (Chandra et al., 2012). Increased susceptibility of the aged brain to ischemic injury arises from reductions in microvascular density (Sonntag et al., 1997), insufficient basal perfusion (Farkas and Luiten, 2001), and impaired vascular responsiveness (Kecskés et al., 2023), in part driven by endothelial dysfunction (Ungvári et al., 2018). Further, reductions in cerebral blood flow (CBF) following vascular occlusion, more pronounced in aged brains, increase the likelihood of spreading depolarization (SD) (Menyhárt et al., 2017; Hertelendy et al., 2017), a key mediator of ischemic injury progression (Hartings et al., 2017). In the aged ischemic brain, SD is characterized by delayed repolarization and impaired CBF responses (Clark et al., 2014; Menyhárt et al., 2015), features that are recognized as biomarkers of exacerbation of neuronal injury (Dreier et al., 2017).

Beyond aging, extracranial carotid stenosis represents a clinically relevant vascular comorbidity that further increases AIS risk (Flaherty et al., 2013), particularly in elderly populations (Sertedaki et al., 2020). Unilateral carotid stenosis contributes to ischemic vulnerability through embolism following plaque destabilization and through chronic hemispheric hypoperfusion resulting from luminal narrowing (Caplan and Hennerici, 1998). Despite its high clinical prevalence, carotid stenosis remains underrepresented in experimental AIS models. Therefore, we incorporated both advanced age and carotid stenosis into our preclinical AIS model to improve translational relevance and to evaluate therapeutic strategies under comorbid conditions that more closely reflect the pathophysiological complexity of ischemic stroke in elderly patients.

The mechanistic basis of age-related cerebrovascular fragility is multifactorial. Emerging evidence implicates chronic cellular senescence and inflammaging as upstream drivers of cerebrovascular vulnerability during aging. Senescent cells accumulate within vascular and perivascular compartments, undergo stable cell-cycle arrest while resisting apoptosis, and acquire a senescence-associated secretory phenotype (SASP) (van Deursen, 2014). Vascular senescence sustains a proinflammatory milieu, promotes endothelial stress and impairs metabolic flexibility (Franceschi et al., 2018). Beyond secreting proinflammatory mediators (including cytokines, chemokines, matrix metalloproteinases, and growth factors), senescent brain endothelial cells exhibit increased expression of adhesion molecules, thereby promoting leukocyte adhesion and inflammatory activation (Real et al., 2024). Moreover, endothelial senescence has been causally linked to age-related cerebral microvascular rarefaction and impaired neurovascular coupling in experimental models of aging (Csik et al., 2025). Collectively, these senescence-associated cerebrovascular maladaptations are predicted to predispose the aged brain to more severe injury following AIS and raise the possibility that senolytic therapies that selectively eliminate senescent cells may mitigate stroke severity and improve outcomes in older individuals (Ouvrier et al., 2024).

Among experimental senolytics, the combination of dasatinib (a tyrosine kinase inhibitor) and quercetin (a plant-derived flavonoid with antioxidant properties) (D+Q), has emerged as a potent senotherapeutic regimen (Al-Naggar et al., 2020), and has advanced to clinical trials for chronic psychiatric and neurodegenerative disorders, including Alzheimer’s disease (Gonzales et al., 2023; Millar et al., 2025). The synergistic activity of D+Q is mediated by the induction of apoptosis in senescent cells; specifically, dasatinib inhibits pro-survival Src family kinases, while quercetin suppresses the anti-apoptotic protein Bcl-xL (Zhu et al., 2015). Despite significant recent interest in senolytics, the effects of these agents in general, and D+Q in particular, on AIS outcomes remain largely unexplored. To address this knowledge gap, we designed the following preclinical study to test whether D+Q could be repurposed as an age-tailored therapeutic to improve stroke outcomes.

Carotid stenosis was induced by unilateral common carotid artery occlusion in aged rats, thereby modeling a high-risk state for subsequent AIS and serving as a diagnostic trigger for initiation of chronic senolytic therapy with D+Q. Two weeks later, the distal segment of the middle cerebral artery was transiently occluded (dMCAO) to model ischemia-reperfusion injury, as occurs in AIS followed by recanalization. Compared with untreated rats, we assessed infarct volume and SDs as biomarkers of injury progression (Varga et al., 2020), CBF dynamics during AIS (Bálint et al., 2019), and senescent cell burden within cerebrovascular compartments (Kecskés et al., 2023). In addition, cerebral angiogenesis and both central and systemic inflammatory responses were characterized using a protein array (Haab, 2006). The same platform was used to evaluate metabolic status by quantifying proteins in blood serum involved in glucose metabolism, in conjunction with blood glucose measurements. Collectively, the acquired preclinical data identify D+Q as a promising age-tailored therapeutic strategy to mitigate AIS severity while improving inflammatory and metabolic profiles in aged animals.

## Materials and Methods

### Experimental groups

All experimental procedures were approved by the National Food Chain Safety and Animal Health Directorate of Csongrád County, Hungary (No. II./761/2024). The study was conducted in accordance with the guidelines of the Scientific Committee of Animal Experimentation of the Hungarian Academy of Sciences (updated Law and Regulations on Animal Protection: 40/2013. (II. 14.) Gov. of Hungary), which aligns with the European Directive 2010/63/EU on the protection of animals used for scientific purposes. The study is reported in compliance with the ARRIVE guidelines (Percie et al., 2020).

Old (18–24 months; 695±95 g) and young (8–10 weeks; 346±77 g) male Wistar rats (n=39) were housed under controlled environmental conditions at a constant temperature of 23 °C with a 12:12-hour light-dark cycle. Standard rodent chow and tap water were provided *ad libitum*. Animals were randomly assigned to one of three experimental groups (n=12 per group): a young, untreated control group; an aged, untreated group; and an aged group treated with dasatinib and quercetin (D+Q). Animals were matched for all other relevant parameters. Sample size determination was based on power analysis to ensure 80% statistical power (β=0.20), using standard deviations of electrophysiological and hemodynamic parameters from previous studies and estimated intergroup differences in mean values. Calculations were performed using SPSS version 20 (Vanderbilt University, USA) and confirmed with GPower 3.1 (Heinrich Heine University, Düsseldorf, Germany).

### Surgical Procedures

Prior to the surgical procedure, blood samples were collected from the tail vein following an overnight fast, and glucose concentrations were determined using an Accu-Chek Active glucometer (Roche Diagnostics GmbH, Mannheim, Germany).

Anesthesia was induced using inhalation of isoflurane (1.5–2%) delivered in a humidified mixture of O_2_ and N_2_O (2:3 ratio), with the animals breathing spontaneously. Lidocaine (10 mg/mL, Egis) was applied to each tissue layer upon incision to provide additional analgesia. To maintain normothermia (37.2 °C), a rectal probe-controlled heating system for small animals (Harvard Apparatus, USA) was used. To suppress airway mucus secretions associated with isoflurane anesthesia, atropine (0.1 mL, 0.1%) was administered intramuscularly. A thin layer of ophthalmic gel (Corneregel, Bausch & Lomb) was applied to the eyes to prevent corneal drying during anesthesia. The surgical preparation followed previously described protocols (Kecskés et al., 2023; Kecskés et al., 2025).

All animals underwent unilateral common carotid artery occlusion (1CCAO) to mimic carotid stenosis. After a midline neck incision, the right common carotid artery was carefully dissected free from the vagus nerve over a 1–2 mm segment and ligated. The skin was closed with sutures, and the surgical site was treated with Betadine (10 mg/mL, Egis) and Lidocaine (10 mg/mL, Egis) for local antisepsis and analgesia. Animals received saline (37 °C) subcutaneously for hydration.

Following 1CCAO, a two-week survival period was implemented. During this time, some aged animals received D+Q (n=12), while control animals (young and aged) received vehicle only (n=12-14/group). Dosages were based on previous studies as follows. Dasatinib (Sigma Aldrich, CDS023389) at 5 mg/kg and quercetin (Sigma Aldrich, Q4951) at 50 mg/kg body weight were administered intraperitoneally (Zhu et al., 2015; Krzystyniak et al., 2022; Zhu et al., 2020). The compounds were dissolved in dimethyl sulfoxide (DMSO) and diluted in physiological saline to a final DMSO concentration of 0.1%. The first dose was administered immediately after 1CCAO, followed by subsequent injections every other day. The final dose was given prior to anesthesia and surgical preparation. No adverse behavioral effects, changes in appetite, fluid intake, or weight loss were observed during D+Q treatment.

After the treatment period, dMCAO was performed on the right side to induce acute ischemic stroke (AIS). The animals were anesthetized again with isoflurane as described above, and placed in a stereotaxic apparatus (Stoelting Co., Wood Dale, IL, USA). After scalp incision, a cranial window was drilled in the right temporal bone (ProLab Basic, Bien Air 810, Switzerland) to occlude the distal branch of the middle cerebral artery and to place an electrode for electrophysiological recording. A second small craniotomy was created in the frontal bone to provide the opportunity to induce spreading depolarizations (SDs) with 1M KCl. The dura was incised in both windows. In addition, the right parietal bone was thinned to allow laser speckle contrast imaging (LSCI) (PeriCam PSI HR, Perimed AB, Sweden) for cerebral blood flow (CBF) monitoring within the ischemic area.

### Experimental Protocol

Once the preparation was completed, a 5-minute baseline recording was taken. Then, AIS was induced by microclip dMCAO for 60 minutes. During this ischemic period, spontaneous SDs were observed and monitored. Reperfusion was achieved by removing the clip, followed by a 60-minute observation period. Subsequently, SDs were evoked by placing a cotton ball soaked in 1M KCl on the cortical surface through the rostral craniotomy left in place for 1 hour, refreshed every 15 minutes, as previously described (Szabó et al., 2017; Szabó et al., 2021). The experimental protocol concluded by transcardial perfusion for immunocytochemistry (n=4/group), or by collecting brain samples for protein array analysis (n=3/group) (Fig. S1).

### Monitoring of Electrophysiological Activity and Cerebral Blood Flow

Electrophysiological recordings (LFP and DC potential) were performed using an Ag/AgCl electrode inserted into a glass capillary filled with physiological saline (120 mM NaCl). The electrode was positioned 700 μm deep in the right somatosensory cortex, referenced to a subcutaneous neck electrode. The tip diameter of the capillary was 20 μm. Signals were amplified using a high-input-resistance preamplifier (NL102GH, NeuroLog System, Digitimer Ltd., UK), connected to a differential amplifier (NL106, NeuroLog System, Digitimer Ltd., United Kingdom), and a signal conditioning and filtering unit (NL125, NL144, NL530, NeuroLog System, Digitimer Ltd., United Kingdom). High-frequency noise from the electrical grid was eliminated using a bandpass filter (HumBug, Quest Scientific Instruments Inc., Canada). Data acquisition was performed at 1 kHz (Labchart, ADInstruments, Australia), and all procedures followed previous descriptions (Szabó et al., 2021; Varga et al., 2020; Kecskés et al., 2023; Kecskés et al., 2025). Electrophysiological data were analyzed using AcqKnowledge 4.2 (Biopac Systems, USA). Parameters were derived from DC-recorded SD waveforms: duration at half amplitude, repolarization slope, and area under the curve (AUC).

Cerebral blood flow (CBF) was monitored by laser speckle contrast imaging (LSCI) (Perimed AB, Sweden). A transparent UV-curing gel (Permabond) was applied to the thinned parietal bone to provide transparency, and the camera was positioned ∼10 cm above the skull to ensure full field coverage. In addition to extracting CBF traces from defined regions of interest (ROIs), spatial CBF distribution across the cortical parenchyma was estimated using CBF maps generated at defined phases of the experimental protocol, as previously described (Clark et al., 2014; Bálint et al., 2019). After removal of motion-related artifacts, the relative area occupied by pixels corresponding to defined perfusion ranges within the region beneath the thinned parietal bone was quantified using custom-made software.

### Immunohistochemistry

Transcardial perfusion was performed under deep isoflurane (3-4%) anesthesia, beginning with physiological saline followed by 4% paraformaldehyde (PFA). Brains were collected and fixed overnight in 4% paraformaldehyde. On the next day, tissue samples were transferred to 30% sucrose in phosphate-buffered saline (PBS) for cryoprotection. Coronal sections (20 μm) were prepared using a freezing microtome (Leica CM 1860 UV, Leica, Germany). Regions for analysis were selected at three rostrocaudal levels based on Paxinos and Watson coordinates: 1 mm rostral to bregma (cortex and striatum), 3 mm and 6 mm caudal (hippocampus dentate gyrus and CA1).

Sections were blocked with 10% normal goat serum (NGS, Merck, USA) to prevent non-specific binding. The sections were labeled with various immunohistochemical markers for microscopic analysis, similar to those previously reported (Kecskes et al., 2023): Anti-p16 (rabbit, Sigma Aldrich, SAB5700620, 1:100, 1 h) for detection of senescent cells; Anti-αSMA (mouse, Genetex, U.S.A., GTX73419, 1 h) for smooth muscle cells; Anti-GFAP (mouse, Sigma Aldrich, Hungary, G3893, 1:1500, 1 h) for astrocytes; Anti-CD31 (mouse, Santa Cruz Biotechnology, sc-376764, 1:50, 1 h) for endothelial cells and Anti-PDGFRβ (mouse, Abcam, ab69506, 1:200, 1 h) for pericytes. Neurons were stained with NeuN antibody, revealing the ischemic lesion (rabbit, Abcam, ab177487, 1:500, 1 h). Co-localization of p16 with other cell-type markers enabled identification of senescent smooth muscle cells, astrocytes, endothelial cells and pericytes.

Secondary antibodies included goat anti-rabbit Alexa Fluor 488 (green) (Thermo Fisher, U.S.A., A32731, 1:100, 1 h) for p16 and NeuN, and goat anti-mouse Alexa Fluor Plus 555 (red) (Thermo Fisher, A32727, 1:100, 1 h) for all other cell types. Sections were mounted with DAPI-containing Fluoromount G (Thermo Fisher, U.S.A., 00-4959-52) and visualized using a Leica DM LB2 fluorescence microscope (Leica Microsystems Wetzlar GmbH., Wetzlar, Germany). Images were captured with a Leica DFC250 color camera. Additional imaging for NeuN and CD31 analysis was conducted on Olympus IX83 inverted microscope integrated with the Olympus ScanR high-content imaging system (Olympus, Tokyo, Japan), equipped with an Orca2 camera (Hamamatsu Photonics K.K., Shizuoka, Japan).

Microscopic images obtained from the contralateral hemisphere were analyzed in triplicate for each animal, and mean values were calculated for subsequent analyses. For NeuN-stained coronal sections, the ischemic lesion was manually delineated in ImageJ, and the enclosed area was measured. Infarct size was calculated for each section as the ratio of the delineated lesion area to the ipsilateral cortical area and expressed as a percentage; values were then averaged per animal. Immunohistochemical images stained for p16, αSMA, GFAP, CD31, and PDGFRβ were evaluated by two independent observers using manual cell counting (40× magnification, 1 mm^2^ fields).

### Protein Array Analysis

To identify cytokines altered by D+Q treatment after stroke, serum and brain samples were collected. Serum was obtained from five rats per group by cardiac puncture, allowed to clot at room temperature for 45 minutes, and centrifuged (4000 rpm, 15 min). Samples were stored at −20 °C until analysis. Brain homogenates (ischemic cerebral cortex) were prepared from three animals per group using RIPA buffer containing protease inhibitors (leupeptin, pepstatin), centrifuged at 13,000 g (15 min), and stored at −80 °C.

Multiplex semiquantitative immunoassays enable efficient profiling of proteomic changes by allowing the simultaneous detection of multiple proteins. Cytokine profiles were assessed using the Proteome Profiler Rat XL Cytokine Array kit (ARY030, R&D Systems, Minneapolis, MN, USA), containing 79 cytokine antibodies in duplicate. Arrays were processed per the manufacturer’s protocol. Positive controls were located in three array corners. Signal intensity was quantified using ImageJ (v1.54g) with the Protein Array Analyzer macro (v1.1.c). Heatmaps were generated using the R statistical environment (R Core Team, 2025, Vienna, Austria; https://www.R-project.org), and cytokine-associated pathways were analyzed using the STRING platform (version 12; https://string-db.org/) based on log_2_ fold-change (log_2_FC) values derived from the protein array dataset.

### Data Analysis

Three aged animals were lost during surgical interventions (one during 1CCAO and two during dMCAO). All surviving animals were included in the data analysis. Blinding of data analysis was implemented by assigning coded identifiers to samples that concealed experimental group allocation. Investigators performing the analyses were unaware of sample identities during analysis. Data were analyzed independently by multiple investigators. Statistical analyses were performed using SigmaPlot 12.5 (Jandel Scientific, USA). Data are presented as mean±standard deviation. Normality was assessed using the Shapiro-Wilk test. Normally distributed data were analyzed by one way ANOVA followed by Holm-Sidak post hoc testing, whereas non-parametric data were analyzed using the Kruskal-Wallis test followed by Dunn’s post hoc analysis. Repeated-measures data were assessed using two-way RM ANOVA with Holm-Sidak post hoc testing. Statistical significance was set at p<0.05* and p<0.01**. Specific statistical tests are indicated in the corresponding figure legends.

## Results

### Dasatinib and quercetin reduce age-related increases in infarct size

In this study, AIS was induced by dMCAO superimposed on prior unilateral 1CCAO. CBF immediately after dMCAO decreased sharply to approximately 64.5-66.7% of the post-1CCAO baseline in both young and aged animals (Fig. 1A-B). In D+Q-treated aged animals, the reduction in CBF was significantly attenuated, reaching approximately 76.7%. Importantly, NeuN immunostaining revealed significantly larger cortical infarcts in aged animals compared to young controls, whereas D+Q treatment reduced infarct size in aged animals toward levels observed in the young (5.07 ± 3.14% vs. 13.01 ± 3.56% vs. 5.72 ± 2.38%; Old D+Q vs. Old vs. Young) (Fig. 1A).

**Figure 1.**
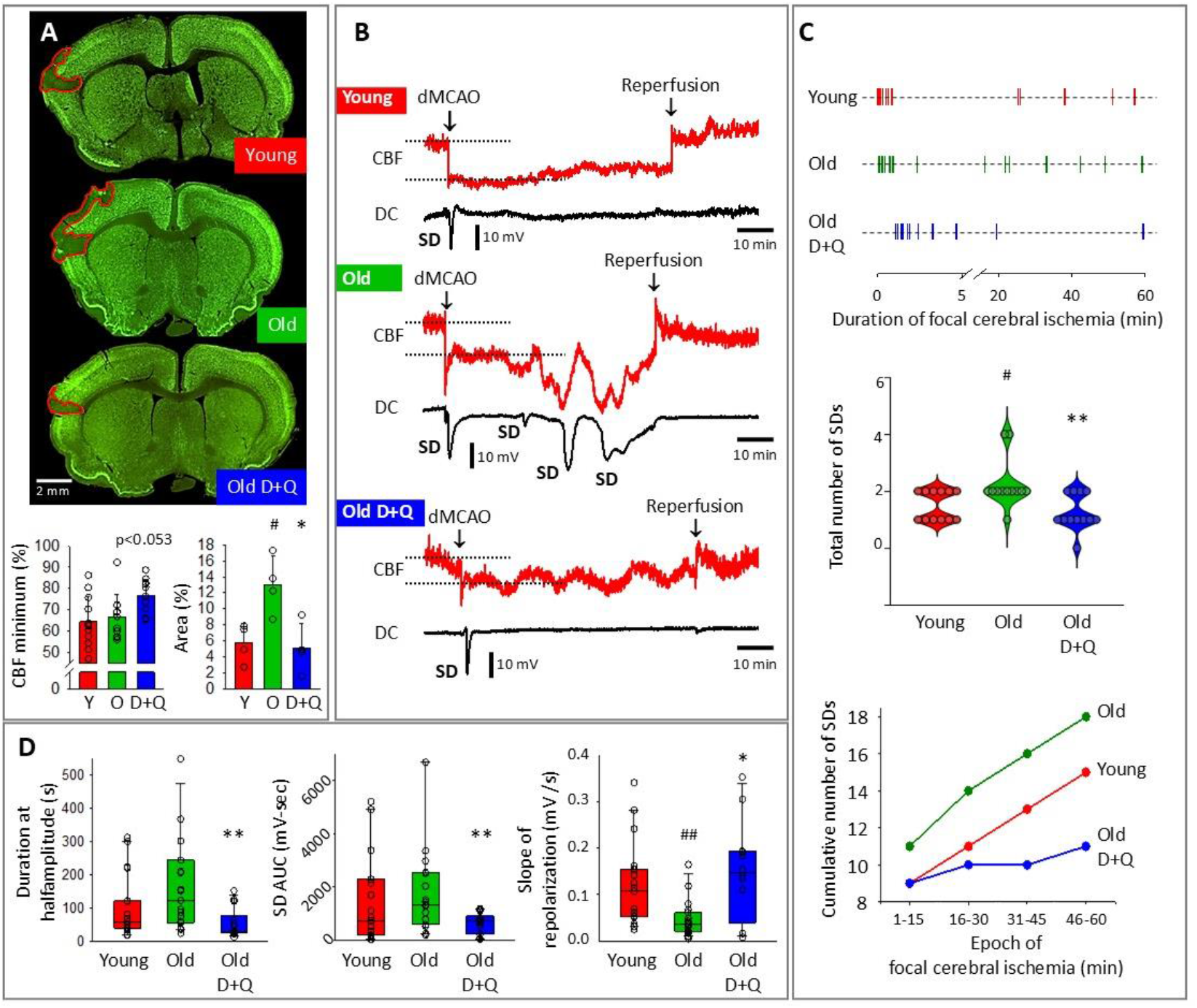
Dasatinib and quercetin (D+Q) reduce infarct size and inhibit spontaneous spreading depolarizations (SDs) during focal cerebral ischemia. **A**, Representative coronal brain sections at the level of the striatum, immunostained for NeuN, show cortical ischemic infarcts 3 h after ischemia induction (outlined in red) in each experimental group. The minimum CBF after dMCAO is expressed relative to the post-1CCAO baseline, and infarct size is expressed as a percentage of the ipsilateral cortical surface area. **B**, Representative recordings of cerebral blood flow (CBF, red) and direct current (DC) potential (black) in each group (Young, Old, Old + D+Q) show CBF reduction and spontaneous SDs during 60 minutes of distal middle cerebral artery occlusion (dMCAO), followed by reperfusion. Spontaneous SDs are indicated by sharp negative DC shifts. **C**, Frequency and temporal distribution of spontaneous SDs during the 60-minute ischemic period across all animals in each experimental group, total number of SDs, and cumulative number of SDs over 15-minute epochs reveals a significant age-related increase in SD incidence, which is mitigated by D+Q treatment. **D**, Quantitative analysis of SD characteristics: area under the curve (AUC), duration at half amplitude, and rate of repolarization. In panel A, data are shown as mean±SD. In panel C, data are presented as violin or scatter plots. In panel D, data are shown as box plots with individual data points overlaid. Normality of distributions was tested using the Shapiro–Wilk method (A, p=0.274 and p=0.692; C, p<0.050; D, p<0.050; p<0.050; p<0.050). For normally distributed data (A), two-way ANOVA followed by Holm–Sidak’s multiple comparisons test was applied. For non-normally distributed data (C-D), statistical analysis was performed using the Kruskal–Wallis test followed by Dunn’s post hoc test. Significance levels: p<0.05^#^ and p<0.01^##^ vs. Young; p<0.05* and p<0.01** vs. Old.

### Dasatinib and quercetin inhibit spontaneous spreading depolarization during cerebral ischemia

SDs are recognized as biomarkers of injury progression in ischemic brain tissue (Dreier et al., 2017). Therefore, we evaluated the impact of D+Q treatment on the pattern of SDs during ischemia. SDs occurred spontaneously after ischemia onset in all groups (Fig. 1B). In young animals, SDs were detected at a relatively low frequency, whereas aged animals exhibited a higher incidence of SDs (1-4 vs. 1-2 events, Old vs. Young). D+Q treatment significantly reduced the frequency of spontaneous SDs in old rats (0-2 vs. 1-4 events, Old D+Q vs. Old), as also reflected by the cumulative number of SDs over 15-minute epochs (Fig. 1C). Aging was associated with delayed recovery from SD, reflected by slower repolarization (0.06±0.09 vs. 0.12±0.09 mV/s, Old vs. Young) and a trend toward longer SD duration. D+Q exerted robust effects by reducing SD duration (54±44 vs. 189±214 s, Old D+Q vs. Old) and AUC (643±385 vs. 2074±2252 mV × s, Old D+Q vs. Old), and by accelerating repolarization (0.15±0.11 vs. 0.06±0.09 mV/s, Old D+Q vs. Old), thereby restoring each parameter toward values observed in young controls (Fig. 1D). These data are in agreement with infarct size estimated in the experimental groups.

### Dasatinib and quercetin do not reverse age-related cerebrovascular dysfunction

An impaired CBF response to SD is thought to delay repolarization following the event (Dreier, 2011). Therefore, we next examined how D+Q treatment affected the CBF response to SD. When a detectable CBF response occurred, its kinetics were either biphasic (an initial transient decrease in CBF followed by hyperemia) or purely hyperemic (only the hyperemic component was observed) (Fig. 2A). As reported previously, responses including an early decrease in CBF were more likely to occur near the ischemic focus and the origin of SD, and gradually transitioned into hyperemic responses as they propagated from the ischemic core to well-perfused tissue in young animals (Fig. 2A-B, from ROI1 to ROI3). This trend was not observed in aged groups, in which biphasic responses remained prevalent even at greater distances from the ischemic core (Fig. 2B). However, the amplitudes of both the initial hypoperfusive component and the subsequent hyperemic component of the CBF response were similar across experimental groups (overall means: -7.9±4.7 and 12.9±11.5%, respectively). D+Q treatment did not result in a statistically significant improvement in the CBF response to SD.

**Figure 2.**
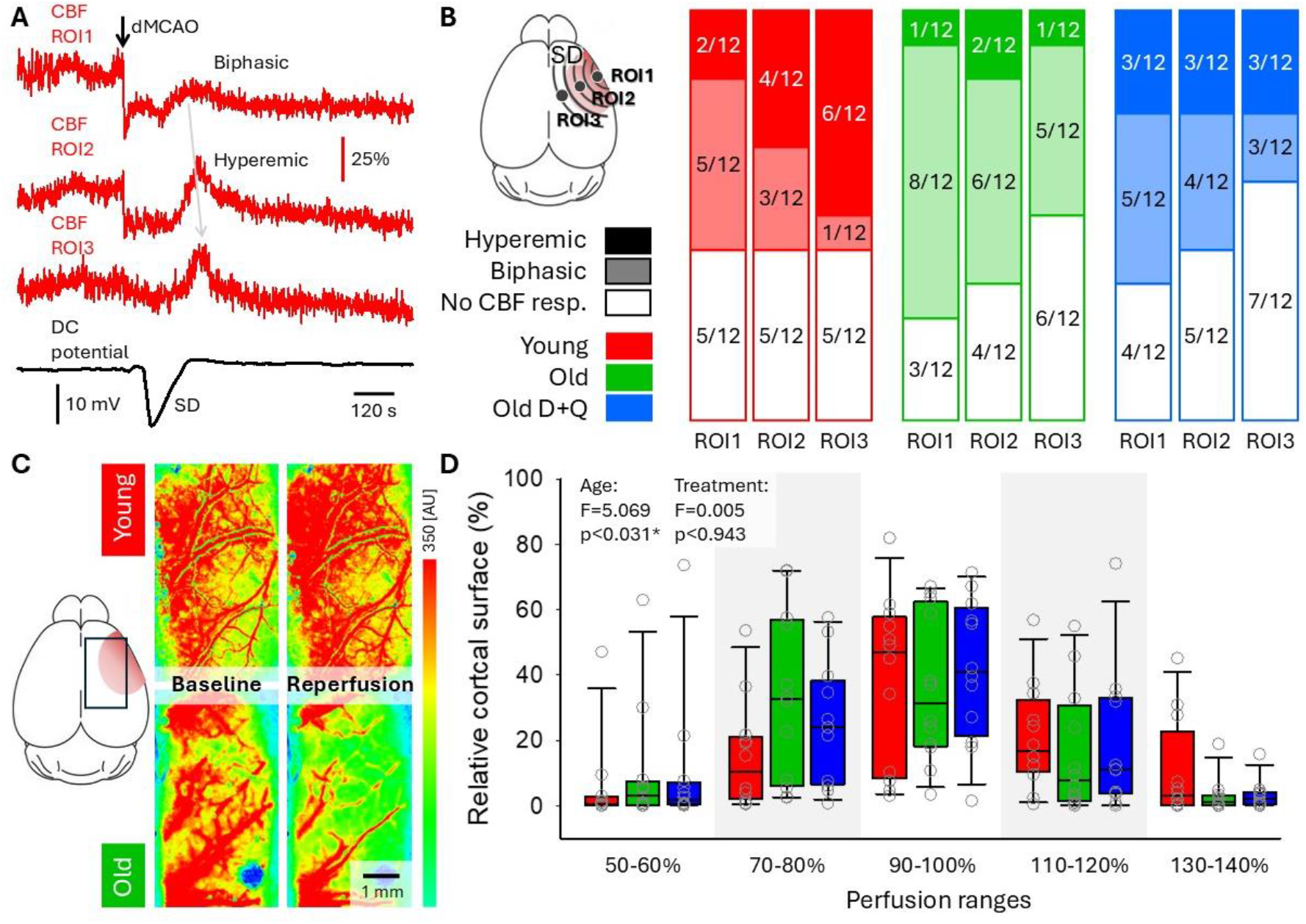
Aging compromises cerebrovascular function, with no effect of dasatinib and quercetin (D+Q). **A**, Representative recordings of cerebral blood flow (CBF, red) from regions of interest (ROIs) across the parietal cortex, along with direct current (DC) potential (black), in a young animal, illustrating the propagation of the CBF response to spreading depolarization (SD) following distal middle cerebral artery occlusion (dMCAO). Note the transition from a biphasic to a hyperemic response along the path of SD. **B**, Incidence of biphasic and hyperemic CBF responses at ROIs located at increasing distances from the ischemic focus across experimental groups. **C**, Cortical perfusion maps obtained an hour after recanalization, demonstrating the relative area adequately reperfused (warm colors) in a young and an aged animal. **D**, Relative cortical area corresponding to predefined perfusion ranges one hour after recanalization. Statistical analysis was performed using a repeated-measures model with two factors (age and treatment). Significance levels: p<0.05* vs. Young.

Finally, the efficacy of reperfusion was evaluated by quantifying the relative cortical area corresponding to predefined perfusion ranges (Fig. 2C-D) (Clark et al., 2014; Bálint et al., 2019). One hour after recanalization, a larger cortical area was adequately reperfused in young animals compared to aged animals, whereas regions with low perfusion were more extensive in aged animals. D+Q treatment in old animals did not result in a statistically relevant improvement.

### Dasatinib and quercetin mitigate age-related cerebrovascular senescence

Cerebrovascular senescence has been shown to correlate with impaired cerebrovascular function (Csik et al., 2025). To investigate the potential protective effects of D+Q treatment against cerebrovascular senescence and to identify the specific cell types involved, we co-localized the senescence marker p16 with endothelial cells (CD31), vascular smooth muscle cells (αSMA), pericytes (PDGFRβ), and astrocytic endfeet (GFAP) (Fig. 3A-D). As expected, senescent cells were more abundant in aged animals than in young controls across all four cell types and brain regions examined (Fig. 3E-H). D+Q treatment consistently reduced the number of senescent cells in all vascular compartments, indicating a broad anti-senescent effect on the neurovascular unit (Fig. 3E-H).

**Figure 3.**
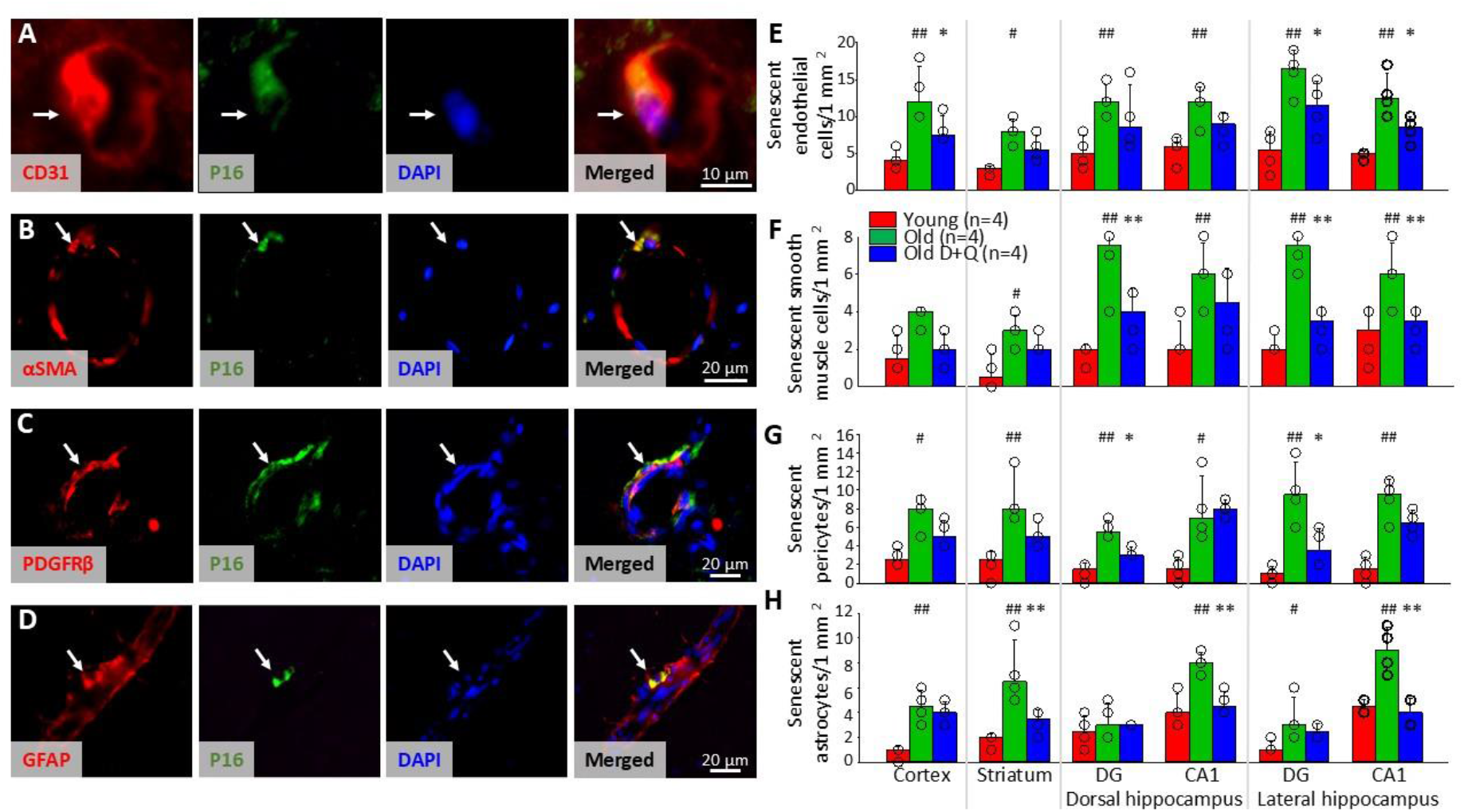
Dasatinib and quercetin (D+Q) mitigate age-related senescence of cerebrovascular endothelial cells, smooth muscle cells, pericytes, and perivascular astrocytes. **A–D**, Representative immunofluorescent images show co-localization of the cellular senescence marker P16 (green) with cell type-specific markers: CD31 for endothelial cells (A), αSMA for vascular smooth muscle cells (B), PDGFR for pericytes (C), and GFAP for astrocytes (D). Cell nuclei are labeled with DAPI (blue). **E–F**, Quantification of P16+ senescent cells co-localized with specific cell markers per 1 mm^2^ across brain regions (cortex, striatum, dentate gyrus [DG], and CA1 region of the dorsal and lateral hippocampus). Data are presented as mean ± SD, with individual data points overlaid. Normality of data distribution was tested using the Shapiro–Wilk test (E, p = 0.432; F, p = 0.335; G, p = 0.191; H, p = 0.078). Subsequently, two-way RM ANOVA followed by Holm–Sidak’s post hoc test was performed. Significance levels: p < 0.05^#^ and p < 0.01^##^ vs. Young; p < 0.05* and p < 0.01** vs. Old.

### Dasatinib and quercetin promote cerebral angiogenesis in aged ischemic brain

Aging is associated with cerebral collateral rarefaction, which worsens stroke outcomes, while ischemia induces compensatory angiogenesis and collateral remodeling (Binder et al., 2024; Faber, 2025). To explore whether senolytic treatment supports vascular remodeling following carotid occlusion, with the potential consequence of improved stroke outcome, we quantified CD31^+^ endothelial cell density across multiple brain regions. Aged animals showed a modest age-related reduction in endothelial density, whereas D+Q treatment significantly enhanced vascular density, particularly in the cortex and striatum (Fig. 4A).

**Figure 4.**
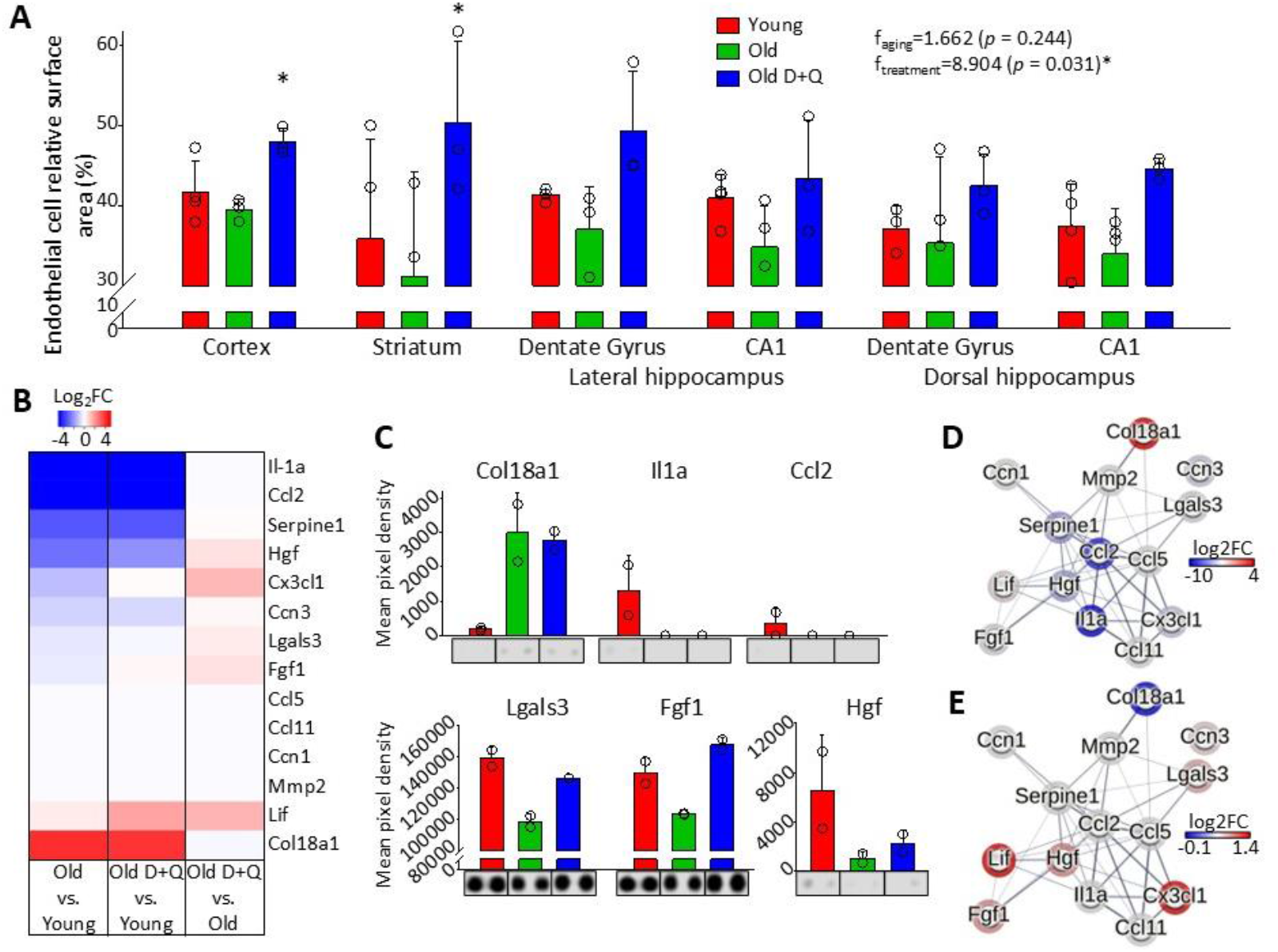
Dasatinib and quercetin (D+Q) enhance cerebral angiogenesis. **A**, Vascular density assessed by the CD31-immunopositive area (endothelial cells) relative to the total area of each brain region (cortex, striatum, dentate gyrus [DG], and CA1 region of the dorsal and lateral hippocampus). **B**, Angiogenesis-related proteins from ischemic brain tissue homogenates, selected from the full protein array and presented as a heatmap, with log2 fold change (log_2_FC) color-coded. **C**, Expression of selected proteins representative of age-related changes without treatment effect (upper row) and proteins restored by D+Q treatment (lower row). Representative array dot images are displayed below each bar graph. **D–E**, STRING network analysis of protein–protein interactions among angiogenesis-related proteins. Line thickness indicates the strength of supporting evidence. Network derived from the Old vs. Young comparison highlights age-associated clustering of angiogenesis-related proteins, reflecting impaired angiogenic signaling in the aged ischemic brain (D). Network derived from the Old + D+Q vs. Old comparison shows treatment-induced normalization of angiogenic signaling pathways, indicating partial restoration of pro-regenerative vascular networks (E). In C, values are expressed as mean ± SD, with individual data points overlaid. Data normality was assessed using the Shapiro–Wilk test (p = 0.235), then statistical analysis was performed using two-way RM ANOVA, followed by Holm–Sidak’s post hoc test. Significance levels: p < 0.05^#^ and p < 0.01^##^ vs. Young; p < 0.05* and p < 0.01** vs. Old.

Protein profiling of brain homogenates revealed broad age-associated alterations in angiogenesis-related proteins (Fig. 4B). The abundance of some vessel-associated proteins increased with age and remained unchanged by D+Q, the most prominent being collagen type XVIII alpha 1 chain (Col18a1), a basement membrane component (Fig. 4C). Conversely, a few proteins were downregulated with age, but their levels were not affected by D+Q; these included interleukin-1α (IL-1a) and C–C motif chemokine ligand 2 (Ccl2) (Fig. 4C). Finally, several pro-angiogenic proteins exhibited age-related reductions that were restored by D+Q, including galectin-3 (Lgals3), fibroblast growth factor 1 (Fgf1), fractalkine (Cx3cl1) and hepatocyte growth factor (Hgf), albeit the latest to a lesser degree (Fig. 4C). STRING interaction analysis demonstrated age-related clustering of impaired angiogenesis-associated proteins (Fig. 4D), whereas D+Q treatment restored connectivity within pro-regenerative vascular networks (Fig. 4E). Together, these findings indicate that D+Q fosters angiogenic signaling and enhances cerebrovascular plasticity after ischemia.

### Dasatinib and quercetin modulate age-related inflammatory protein changes

Senescent cells typically acquire a pro-inflammatory senescence-associated secretory phenotype (SASP). Senolytic agents that eliminate senescent cells are therefore expected to attenuate the pro-inflammatory environment associated with aging and stroke. To assess the impact of D+Q on inflammation, we performed a protein array analysis focused on inflammatory signatures.

Heatmap analysis of brain homogenates revealed a robust age-associated pro-inflammatory shift, which was partially normalized by D+Q treatment (Fig. 5A–B). Analysis of selected protein levels showed age-related upregulation of C–C motif chemokine ligand 21 (Ccl21) and downregulation of fractalkine (Cx3cl1) and vascular cell adhesion molecule 1 (Vcam1) – all of which were markedly counterbalanced by D+Q. Further, intercellular adhesion molecule 1 (Icam1) levels were elevated by D+Q, independent of age (Fig. 5B).

**Figure 5.**
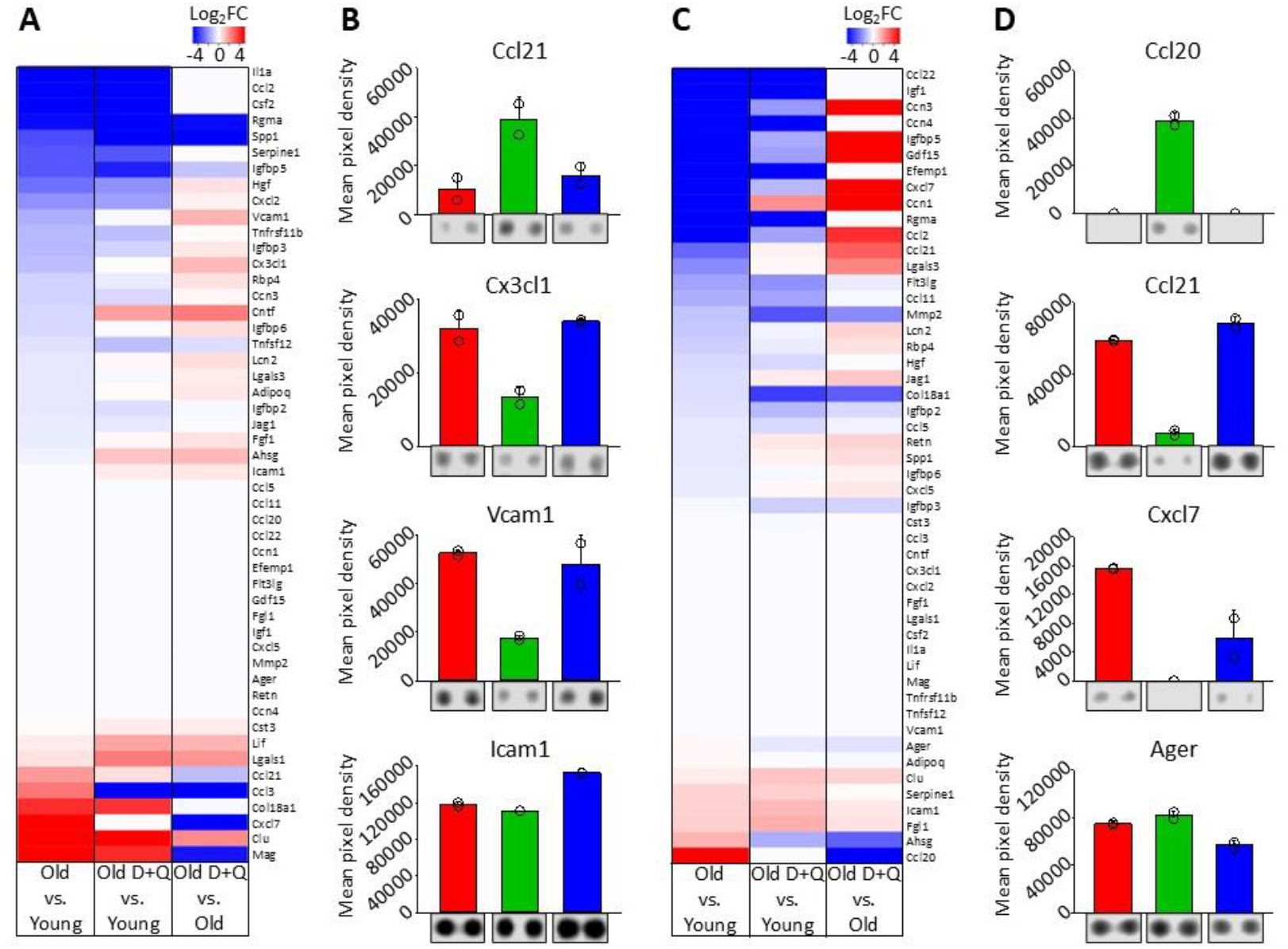
Dasatinib and quercetin (D+Q) modulate the expression of inflammatory protein markers. **A-B**, Proteins involved in inflammation from ischemic brain tissue homogenates. Heatmap showing log_2_ fold change (log_2_FC) values, color-coded (A). Expression of selected proteins representative of the effect of D+Q treatment, with representative array dot images shown below each bar graph (B). **C-D**, Proteins involved in inflammation from blood serum samples. Heatmap showing log_2_ fold change (log_2_FC) values, color-coded (C). Expression of selected proteins representative of the effect of D+Q treatment, with representative array dot images shown below each bar graph (D).

Serum protein profiling further demonstrated systemic inflammatory alterations with aging, including upregulation of C–C motif chemokine ligand 20 (Ccl20) and downregulation of Ccl21 and C– X–C motif chemokine ligand 7 (Cxcl7), all of which were reversed by D+Q. In addition, a few proteins, including RAGE (Ager), exhibited treatment-related reductions without age-dependent changes (Fig. 5E–F). Together, these findings support both central and peripheral immune modulatory effects of D+Q treatment.

### Dasatinib and quercetin improve glucose homeostasis and metabolic protein networks

Hyperglycemia impairs cardiovascular function and exacerbates ischemic brain injury, so we measured blood glucose concentrations in the experimental animals. Significantly elevated glucose levels were detected in aged compared to young animals in our cohort, and this increase was fully reversed by D+Q treatment (7.01 ± 1.08 vs. 9.13 ± 1.11 vs. 7.24 ± 1.99 mmol/L; Old D+Q vs. Old vs. Young) (Fig. 6A).

**Figure 6.**
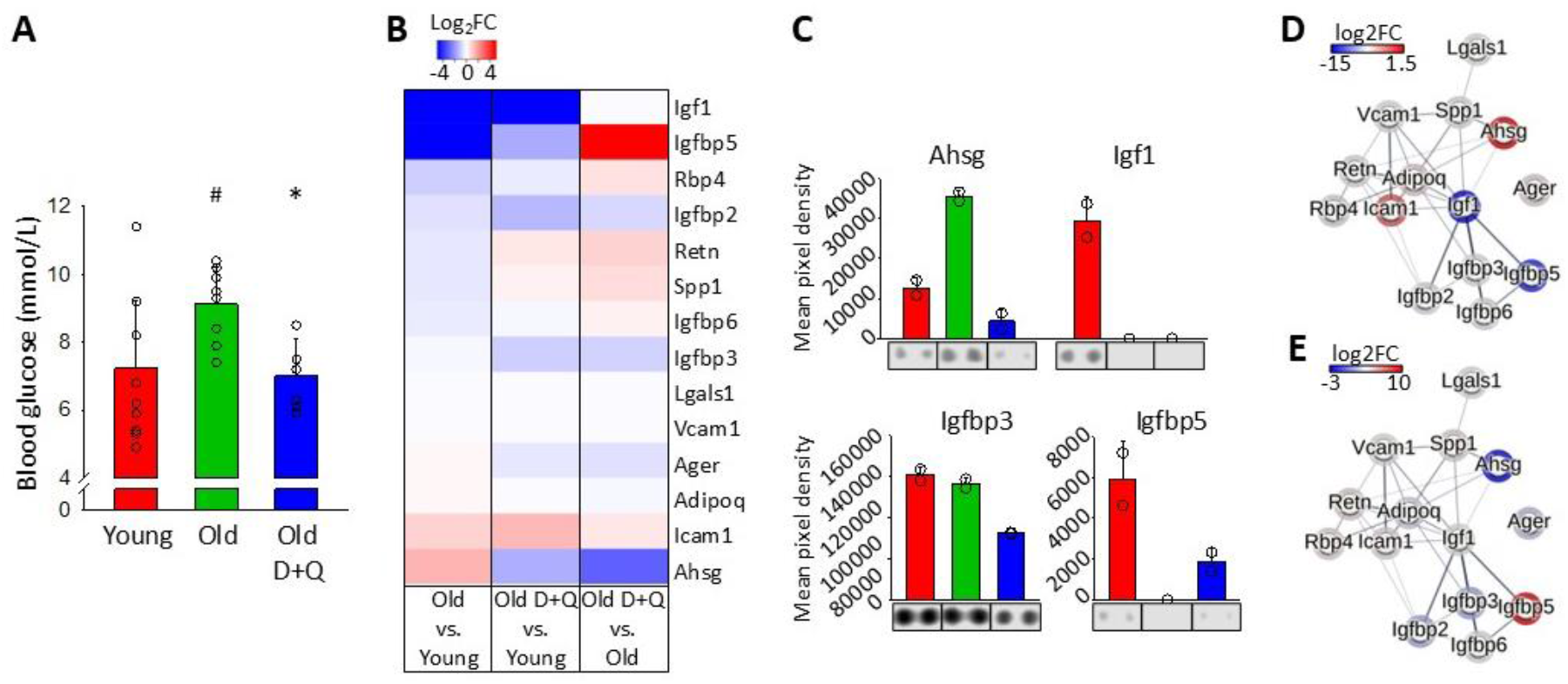
Dasatinib and quercetin (D+Q) lower age-associated increases in blood glucose levels and modulate the expression of glucose-regulatory proteins in rat serum. **A**, Blood fasting glucose concentrations increased with age, an effect counteracted by D+Q. **B**, Changes in the expression of proteins from blood serum, related to glucose metabolism and blood glucose serum levels. Proteins were selected from the full protein array to be presented as a heatmap, with log2 fold change (log_2_FC) color-coded. **C**, Expression of selected proteins representative of aging effects and D+Q treatment. **D-E**, STRING network analysis of protein–protein interactions among proteins related to glucose homeostasis. Line thickness indicates the strength of supporting evidence. Network derived from the Old vs. Young comparison highlights age-associated alterations in the glucose-regulatory network (D). Network derived from the Old + D+Q vs. Old comparison shows treatment-induced partial restoration of glucose-regulatory pathways (E). In A, values are expressed as mean ± SD, with individual data points overlaid. Data normality was assessed with the Shapiro–Wilk test (p=0.188). Group differences were analyzed using one-way ANOVA followed by Holm–Sidak’s post hoc test. Significance levels: p < 0.05^#^ vs. Young; p < 0.05* vs. Old.

In the full serum protein array, several proteins implicated in glucose homeostasis were identified, and heatmap analysis of these proteins revealed age-associated dysregulation of glucose-regulatory pathways (Fig. 6B). Targeted validation showed that D+Q treatment restored age-related increases in some proteins, such as fetuin-A (Ahsg), which is implicated in insulin resistance, and D+Q compensated for age-related decreases in others, such as insulin-like growth factor-binding protein 5 (Igfbp5), a modulator of insulin-like growth factor activity. In contrast, some proteins, including insulin-like growth factor 1 (Igf1), exhibited age-dependent reductions without treatment effects.

Finally, proteins such as insulin-like growth factor-binding protein 3 (Igfbp3) were selectively responsive to D+Q, independent of aging (Fig. 6C). STRING network analysis highlighted age-related disruption of glucose-regulatory protein interactions (Fig. 6D), which was partially normalized by D+Q (Fig. 6E). Together, these findings demonstrate that D+Q alleviates age-associated metabolic dysfunction, with potential implications for improved cerebrovascular function.

## Discussion

In this study, a senolytic cocktail of D+Q was administered guided by carotid artery occlusion as a preventive measure to mitigate the consequences of subsequent AIS. This model reflects a clinically relevant scenario, as carotid stenosis carries a high risk of subsequent AIS and presents an opportunity for preventive intervention. D+Q exert senolytic activity in part by inhibiting pro-survival Src family kinases and suppressing the anti-apoptotic protein Bcl-xL (Zhu et al., 2015). This combination provides a broad-spectrum senolytic effect, as D+Q act synergistically to target different senescent cell survival pathways, making the treatment more effective than either drug alone (Kirkland and Tchkonia, 2020). Importantly, this cocktail selectively induces apoptosis in senescent cells while sparing healthy proliferating and quiescent cells (Kirkland and Tchkonia, 2020). Beyond its senolytic role, quercetin also functions as a potent antioxidant flavonol that suppresses endothelial inflammatory signaling and protects cerebrovascular endothelial integrity (Lu et al., 2026).

Protein kinases – including Src family kinases targeted by D+Q – and their complex interactions have recently been highlighted as druggable targets for the treatment of neurodegenerative disorders (Wu et al., 2025). In a model of acute brain injury, we show that D+Q limited infarct size after AIS in the aged brain (Fig. 1), consistent with the effects of Src kinase inhibitors SKI-606 and SKS-927, which reduced infarct volume and facilitated functional recovery, likely through regulation of cerebrovascular permeability (Liang et al., 2009). We further demonstrate that D+Q inhibited SD (Fig. 1), an electrophysiological biomarker and likely promotor of ischemic injury (Hartings et al., 2017). Intriguingly, Src family kinase activation has been identified as a signaling mechanism that increases the susceptibility of the cerebral cortex to SD (Bu et al., 2018), suggesting a potential mechanistic target through which D+Q may limit SD, as observed here. Finally, aging was characterized by a pro-inflammatory shift in ischemic brain tissue, which was counterbalanced by D+Q (Fig. 5). In particular, the age-related increase in Ccl21 together with a decrease in fractalkine (Cx3cl1) levels is considered indicative of impaired neuron-to-glia communication, leading to microglial activation and pro-inflammatory polarization (de Jong et al., 2005; Bachstetter et al., 2011). The beneficial effects of D+Q on these parameters observed in our experiments are supported by previous complementary findings showing that D+Q attenuated microglial activation, and suppressed neuroinflammation in aged mice (Ogrodnik et al., 2021). Overall, the growing body of evidence implicating Src kinases in stroke pathophysiology and aging aligns well with our findings on the impact of D+Q on tissue injury, SD, and neuroinflammation.

The effects of D+Q on cerebrovascular structure and function were in the focus of our experiments, as the more severe outcome of AIS with aging has been attributed to reductions in microvascular density (Sonntag et al., 1997), insufficient basal perfusion (Farkas and Luiten, 2001), and impaired vascular responsiveness (Kecskés et al., 2023). At the structural level, D+Q eliminated senescent endothelial and mural cells in the cerebral microvasculature (Fig. 4) and promoted cerebral angiogenesis (Fig. 5). The increased levels of Fgf1, Hgf, and Galectin-3 (Fig. 4) indicate systemic rejuvenation of the vascular system (Markowska et al., 2010; Chen et al., 2023). Fgf1 and Hgf deliver strong mitogenic signals to endothelial cells, while Galectin-3 and adhesion molecules such as Icam-1 and Vcam-1 (Fig. 5) support the structural framework and migration needed for vessel formation in various tissues (Koch et al., 1995; Gho et al., 1999). Importantly, senolytics have been shown to modulate angiogenesis in a context-dependent manner. In support of our findings, senolytic therapy targeting endothelial senescence in aged mice improved cerebrovascular density (Csik et al., 2025). Further, the adhesion and integration of endothelial progenitor cells to support vessel growth, which are impaired by aging, were enhanced by D+Q (Lam et al., 2024). Our study demonstrates a restorative, pro-angiogenic effect of D+Q in the aging ischemic brain.

At the functional level, however, D+Q did not improve the CBF response to SD during focal ischemia or the efficacy of reperfusion within the first 60 min after recanalization (Fig. 2), despite previous reports associating senolytic interventions with improved cerebrovascular function (Csik et al., 2025). A limitation of our study is that variation in perfusion following carotid artery occlusion was not monitored, and CBF measurements during dMCAO-induced focal ischemia/reperfusion were expressed relative to the pre-dMCAO baseline. Indeed, the cerebral vasculature upstream of the occluded carotid artery was likely dilated to compensate for the reduced blood supply, and the extent of this compensation was expected to vary across experimental conditions, possibly resulting in different pre-dMCAO baselines. Together, these factors could have masked potential D+Q-induced improvements in cerebrovascular function.

The complex pattern of proteins implicated in brain tissue and systemic inflammation collectively points to D+Q-mediated immune-cell recruitment. Serum levels of Ccl21 and Cxcl7 (also known as neutrophil-activating peptide 2, NAP-2) (Fig. 5C-D), which promote leukocyte recruitment to the central nervous system (Gracia et al., 1994; Engelhardt, 2006), decreased with aging and were restored by D+Q. Likewise, Vcam1, which facilitates endothelial leukocyte adhesion in AIS (Hoyte et al., 2010), was reduced in the aged ischemic brain homogenates and was restored by D+Q (Fig. 5A-B). This effect of D+Q appears counterintuitive, as immune cell recruitment is generally considered pro-inflammatory associated with worse outcomes after AIS. However, in the context of aging, and given the reduction in infarct size observed with D+Q in this study, these changes may instead reflect partial reversal of immunosenescence (Rodrigues et al., 2021) and recovery of a more effective post-ischemic inflammatory response that supports tissue repair.

Finally, we also observed that D+Q reduced blood glucose levels and counteracted age-associated metabolic dysfunction. This is consistent with previous work showing that D+Q decreases adipose tissue inflammation and improves systemic metabolic function in aged mice (Islam et al., 2023), and enhances insulin sensitivity and glucose tolerance in obese mice (Sierra-Ramirez et al., 2020). Given the profound impact of metabolic status on cerebrovascular function (Coucha et al., 2018), improved metabolic homeostasis may have contributed indirectly to the reduced ischemic damage observed in D+Q-treated animals.

In conclusion, our findings demonstrate that senolytic D+Q therapy administered after carotid artery occlusion confers multifaceted protection against subsequent AIS in the aged brain, reducing infarct burden, suppressing SD, alleviating cerebrovascular senescence, and normalizing age-related inflammatory and metabolic disturbances. By targeting fundamental aging mechanisms that exacerbate brain vulnerability to AIS, D+Q enhances the resilience of the aging neurovascular niche. Although additional work is needed to define long-term outcomes and translational feasibility, these results identify senolytic therapy as a promising preventive personalized approach to mitigate the disproportionate impact of AIS in older individuals and warrant further investigation.

